# Tracing Early Migratory Neurons in the Developing Nose Using Contactin-2 (Cntn2)CreERT2

**DOI:** 10.1101/2025.03.27.645843

**Authors:** Enrico Amato, Alexis Semonv, Paolo E Forni

## Abstract

Neuronal migration during embryonic development is a fundamental process. In the developing nose of rodents, neurons that form during early neurogenic waves in the olfactory placode leave this structure to migrate toward or into the developing brain as part of the migratory mass. This mass includes Gonadotropin-releasing hormone-1 (GnRH-1) neurons, pioneer/terminal nerve (TN) neurons, as well as neural crest-derived olfactory glial cells called olfactory ensheathing cells.

There have been a limited number of molecular markers available to effectively trace and functionally manipulate the early migratory neurons that originate in the olfactory region. Contactin-2 (Cntn2), also known as transiently expressed axonal surface glycoprotein-1 (TAG-1), has been used to label various developing neuronal populations, including the commissural neurons of the spinal cord, motor neurons, and TN neurons. Previous single-cell RNA sequencing analyses of the developing olfactory system have identified Cntn2 expression in the TN, suggesting that Cntn2 is a suitable molecular marker for studying nasal migratory neurons. To trace Cntn2 expression in the developing olfactory system, we generated an inducible Cntn2CreERT2 mouse line. In this study, we outline how this mouse line can serve as an effective tool for time-controlled chimeric manipulation of specific neuronal populations of interest.

## Introduction

Contactin-2 (Cntn2), also known as transiently expressed axonal surface glycoprotein-1 (Tag1), belongs to the immunoglobulin (Ig) superfamily (Furley *et al*., 1990). Cntn2 has been implicated in various developmental processes such as cell adhesion (Gurung *et al*., 2018), axonal coalescence (Wolman *et al*., 2008), neurite outgrowth, axon pathfinding, and regulation of neuronal migration (Suter and Jaworski, 2019) .

Cntn2 can facilitate cell-cell contacts through interactions with other cell adhesion molecules and extracellular matrix molecules (Fan *et al*., 2024).

Besides its specific biological functions, Cntn2 has been widely used as a developmental stage-specific molecular marker for various neuronal populations, including commissural neurons of the spinal cord, motor neurons, and putative terminal nerve (TN) neurons in the nose of rodents (Amato *et al*., 2024; Dodd *et al*., 1988; Taroc *et al*., 2017; Tessier-Lavigne *et al*., 1988; Yamamoto *et al*., 1986).

The GnRH-1 neurons are a hypothalamic neuronal population that produces and releases the decapeptide GnRH-1 (Schwanzel-Fukuda and Pfaff, 1989; Wray, 2001; Wray *et al*., 1989). The pulsatile release of GnRH-1 regulates the hypothalamic-pituitary-gonadal hormonal axis, which is responsible for reproductive development and hormone regulation in adulthood (Kaprara and Huhtaniemi, 2018; Pohl and Knobil, 1982; Schwanzel-Fukuda *et al*., 1989).

During embryonic development, the GnRH-1 neurons migrate from the olfactory pit to the hypothalamus as part of a migratory mass that also includes pioneer/TN neurons and olfactory glial cells called olfactory ensheathing cells (Barraud *et al*., 2010; Barraud *et al*., 2013; Forni *et al*., 2011b; Taroc *et al*., 2020b).

The migration of GnRH-1 neurons has been a subject of investigation for several decades (Duittoz *et al*., 2021; Schwarting *et al*., 2007). However, the lack of molecular markers and precise genetic entry points to study, follow and manipulate the early neurons forming in the olfactory area has been a significant limitation.

While for decades, this migration was described to occur along the axons of vomeronasal or olfactory neurons (Schwarting *et al*., 2001; Yoshida *et al*., 1995), other studies suggested that the GnRH-1 neurons invade the brain on the axons of pioneer neurons, forming the putative TN (Amato *et al*., 2024; Demski and Schwanzel-Fukuda, 1987; Schwanzel-Fukuda *et al*., 1987; Schwanzel-Fukuda and Pfaff, 1989; Taroc *et al*., 2020a; Taroc *et al*., 2017).

Peripherin (Jennes, 1992; Wray *et al*., 1994) and Cntn2 (Schwarting *et al*., 2001; Schwarting *et al*., 2004) have been used as markers to follow the development of the GnRH-1 migratory track (Schwarting *et al*., 2001). Cntn2 immunoreactivity is notably stronger in axons than in cell bodies making it challenging to identify which cells express it. However, the question of which cells in the migratory mass express Cntn2 (Taroc *et al*., 2017) has never been clearly addressed because of the transient expression of this gene and the intermingling of axons and cell bodies within the migratory mass (Chao *et al*., 2009).

The migratory neurons that form in the olfactory placode are among the first to express neuronal markers (Fornaro *et al*., 2007; Gong and Shipley, 1995). In a recent study, Prokr2, a gene associated with defective GnRH-1 migration to the brain, and the Microtubule-associated protein 2 (MAP2) have been respectively identified as an early genetic marker and antigen for the presumptive TN neurons in the developing nose (Amato *et al*., 2024). Notably, Prokr2Cre tracing allowed for the identification of Prokr2-expressing TN cells in the early developing nose as a distinct cell population, separate from GnRH-1 immunoreactive neurons and early olfactory and vomeronasal sensory neurons. Single-cell sequencing of the Prokr2-expressing TN cells indicated that these cells are enriched for Peripherin and Cntn2/Tag-1.

Cntn2 loss of function can compromise the development of the nervous system (Suter *et al*., 2020). So, to further characterize the TN and test if Cntn2 can be used as a timely controlled genetic entry point in the developing GnRH-1-TN system, we designed a new line of inducible Cntn2^CreERT2^ mice. The P2A auto cleaving strategy was adopted to faithfully drive Cre expression, relying on endogenous cis-regulatory elements (Fig. 1A), without interfering with Cntn2 endogenous expression and function (Donnelly *et al*., 2001a; Donnelly *et al*., 2001b; Tang *et al*., 2009).

**Figure 1.**
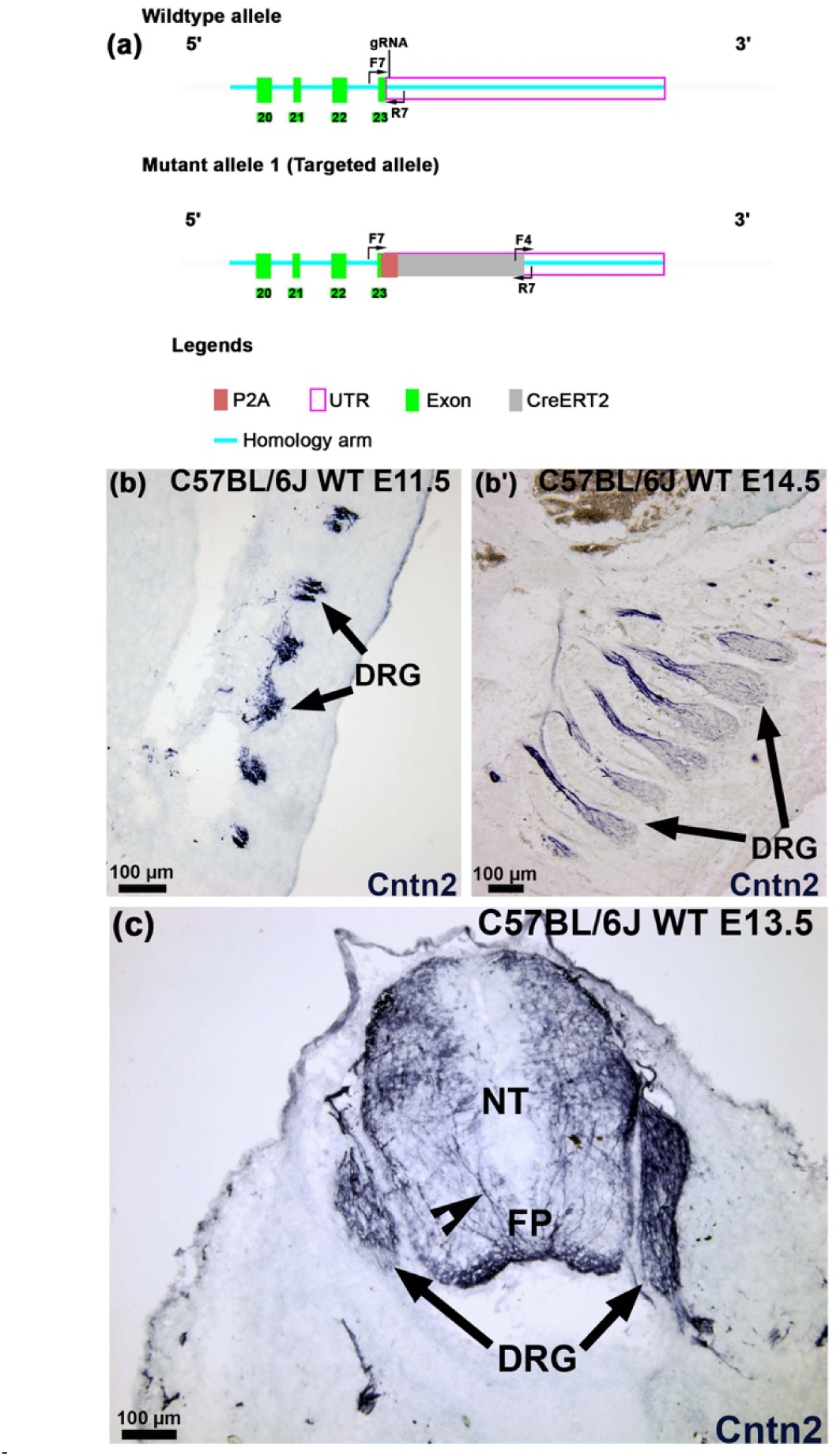
Strategy for generation of Cntn2^CreERT2^ mice and expression of Cntn2 in the Spinal Cord. (a) Construct used in the creation of Cntn2^CreERT2^ mice. Based on strategy report provided by Cyagen. (b-b’) Parasagittal sections of E11.5 and E14.5 C57BL/6J WT embryos immunostained for Cntn2, highlighting expression in the dorsal root ganglia (DRG). (c) Transverse section of an E13.5 embryo, Cntn2 is expressed by the DRG and within the commissural neurons (notched arrowhead) passing through the floorplate (FP) of the neural tube (NT). Scale bars in b–b’, c,100 µm.

In this article, we describe how this new Cntn2^CreERT2^ mouse line can serve as a valuable tool for chimeric manipulation and neuron tracing in a timely and controlled fashion.

## Results

### Cntn2^CreERT2^ recombination faithfully recapitulates Cntn2 expression in the spinal cord and Dorsal Root Ganglia

During development, Cntn2 is transiently expressed by subsets of spinal cord commissural neurons and by neurons in the dorsal root ganglia (Furley *et al*., 1990; Karagogeos *et al*., 1991) (Fig. 1B, C). We tested Cntn2^CreERT2^ mediated recombination by injecting tamoxifen at E11.5 and collecting Cntn2^CreERT2^ ^+/-^/R26tdTomato; Ai14^+/-^ embryos one day post injection (1 DPI) at E12.5 (Fig. 2B). Cntn2 is known to have a transient expression pattern. Immunofluorescent staining against Cntn2 and the reporter gene tdTomato revealed a faithful expression pattern of the reporter gene, overlapping with Cntn2 immunoreactivity in 75% of all the cells that underwent tamoxifen induced Cre recombination 1 day before (Fig. 2C, D). We observed that Cntn2 tracing is specific to areas in the spinal cord where Cntn2 expression is found (commissural neurons and dorsal root ganglia), this shows that the Cntn2^CreERT2^ mouse model accurately traces cells that express Cntn2.

**Figure 2.**
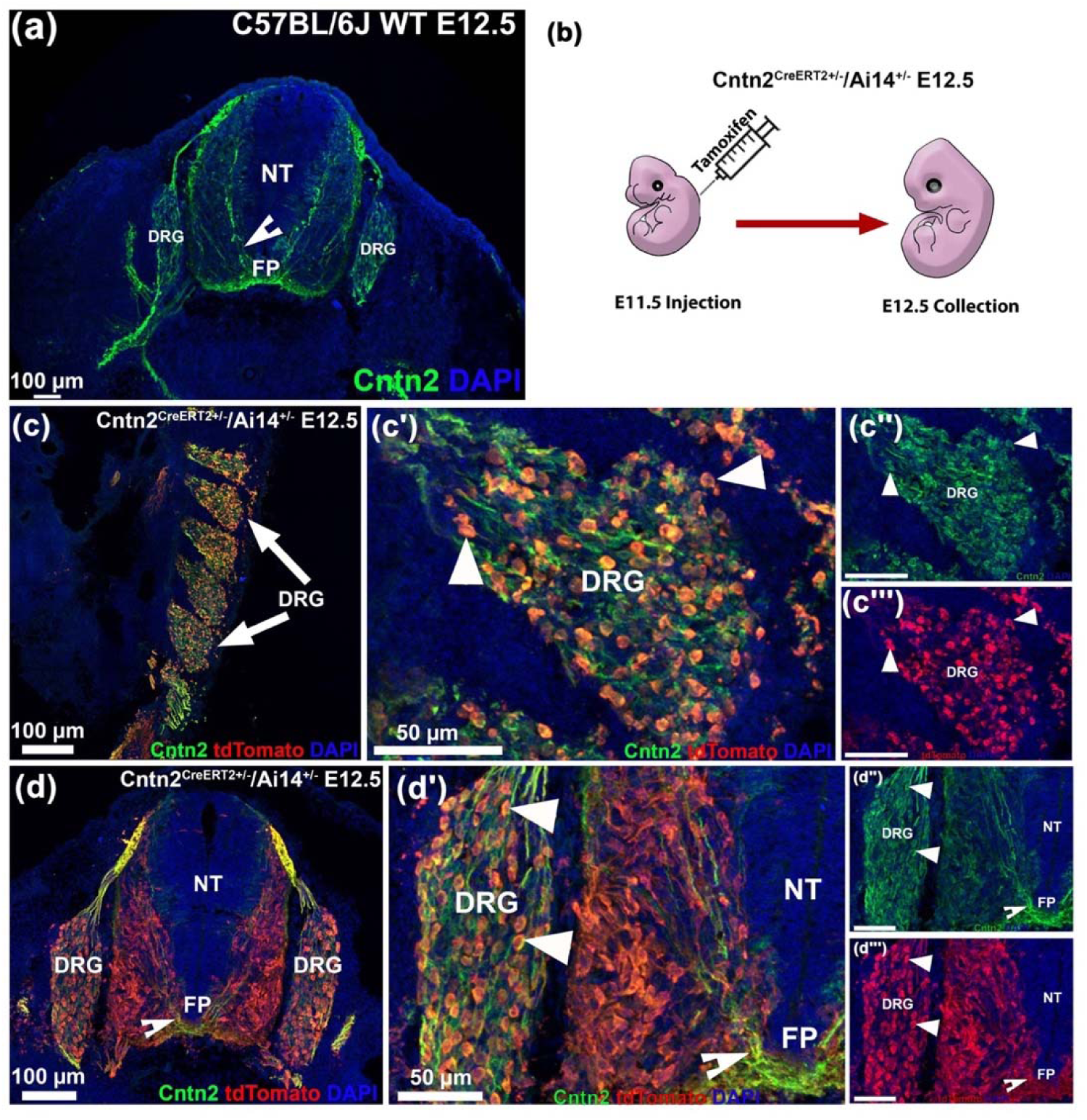
Cntn2^CreERT2^ recombination in the spinal cord overlaps with Cntn2 immunoreactivity. (a) Immunofluorescence for Cntn2 (Contactin 2, green) on transverse E12.5 C57 BL/6J WT tissue reveals the dorsal root ganglion (DRG) proximal to the neural tube (NT) are positive. The commissural neurons are highlighted by Cntn2 immunostaining, passing through the floor plate (FP) (notched arrowhead). (b) Cntn2^CreERT2^ ^+/-^/Ai14^+/-^ mice were injected with tamoxifen at E11.5 and collected at E12.5. (c-c’’, d-d’’) Double Immunofluorescent staining against Cntn2 (green) and tdTomato (red) on parasagittal and transverse E12.5 Cntn2^CreERT2^ ^+/-^/Ai14^+/-^ animals highlights traced neurons immunopositive for Cntn2 in the DRGs. Scale bars in a,c,d 100 µm; c’-c’’’, d’-d’’’, 50 µm.

### Cntn2 is expressed by the neurons of the nasal migratory mass

Cntn2 has been extensively used to follow the trajectory of migrating GnRH-1 neurons (Duittoz *et al*., 2021; Schwarting *et al*., 2001; Taroc *et al*., 2017; Yoshida *et al*., 1995).

In a recent study, we traced the pioneer/TN neurons forming from the olfactory placode, showing that these cells are genetically distinct from the olfactory, vomeronasal, and GnRH-1 neurons (Amato *et al*., 2024). Gene expression profiling of the putative TN cells highlighted the enrichment of Peripherin, Robo3, and Cntn2 expression, which have been previously suggested to mark the GnRH-1 migratory scaffold (Taroc *et al*., 2019; Taroc *et al*., 2017) (Fig. 5A).

Immunohistochemistry against Cntn2 on control animals from E10.5-E13.5 confirmed the Cntn2 immune detectability in axons and in sparse cell bodies in the developing olfactory area (Fig. 3).

**Figure 3.**
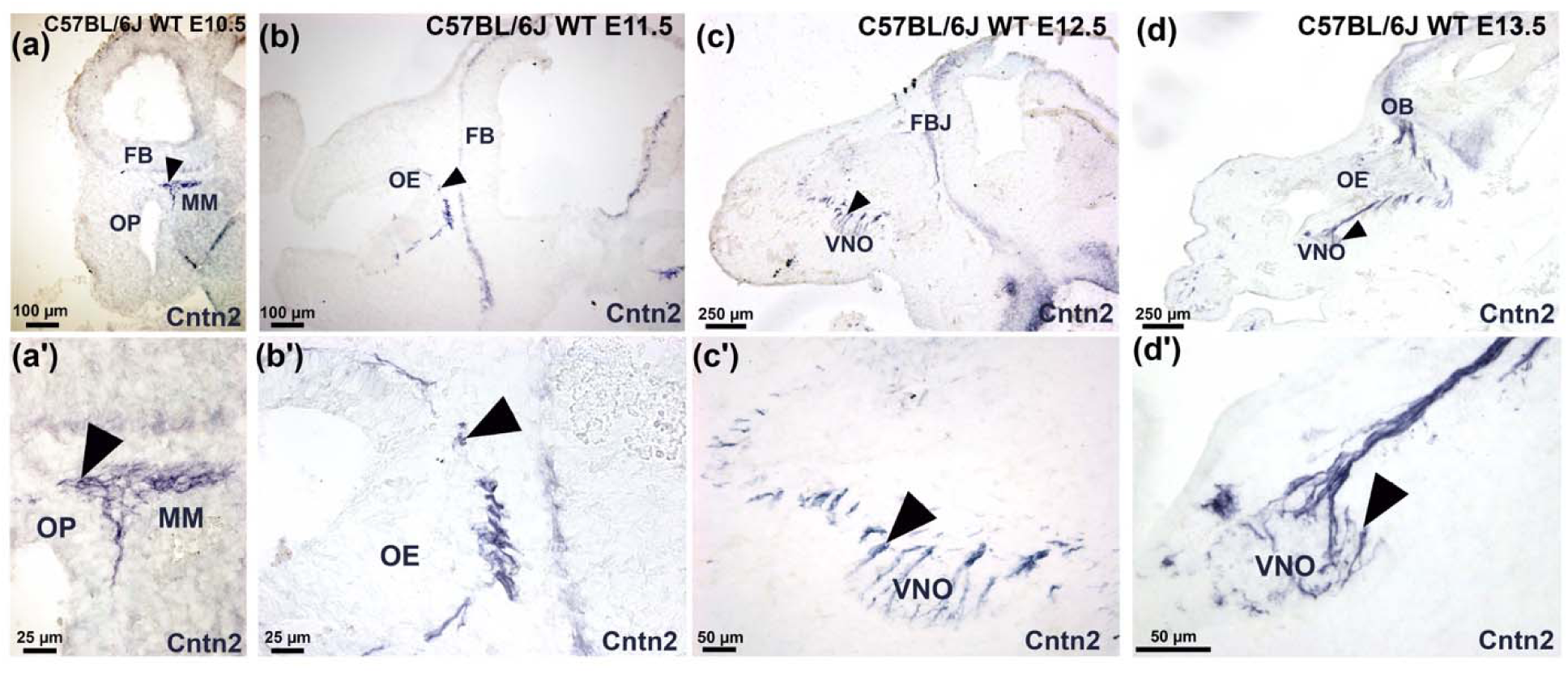
Contactin-2 (Cntn2) is expressed by neurons of the nasal migratory mass. (a-d) developmental time-course of immunohistochemistry against Cntn2 on C57BL/6J WT mice. (a-a’) At E10.5, Cntn2 expression is shown in the migratory pioneer/TN neurons within the migratory mass (MM) emerging from the olfactory placode (OP) and proximal to the developing forebrain (FB) indicated by the black arrows. (b-b’) Cntn2 expression at E11.5 is still present in the pioneer/TN neurons migrating away from the olfactory epithelium (OE). (c-c’) At E12.5, Cntn2 expression is seen in a select group of neurons, labelling cell bodies and fibers outside of the vomeronasal organ (VNO) and adjacent to the forebrain junction (FBJ). (d-d’) Observing Cntn2 expression at E13.5 showing neurons leaving the VNO and projecting toward the brain passing the olfactory bulb (OB). Scale bars in a’, b’, 25 µm; c’, d’, 50 µm; a, b, 100 µm; c, d, 250 µm.

Cntn2 expression was found in migratory pioneer/TN neurons at all stages of development analyzed. At E12.5 and E13.5, Cntn2+ cell bodies and fibers could be observed both outside the VNO and proximal to the forebrain junction (FBJ) (Fig. 3C-D). Previous birth-dating experiments suggested that most of the neurons that become postmitotic between E10.5 and E11.5 migrate out of the developing olfactory system (Fornaro *et al*., 2003; Forni *et al*., 2011a; Taroc *et al*., 2020b; Wray and Hoffman, 1986). Suggesting that the pioneer/TN neurons, like the GnRH-1 neurons, are among the very first neurons to form in the olfactory area.

By analyzing GnRH-1 and Cntn2 immunoreactivity in the developing olfactory system at E11.5, E12.5, and E13.5, we observed very sparse Cntn2 immunoreactivity across the cells in the putative vomeronasal area (Fig. 4), where both GnRH-1 and TN neurons form (Amato *et al*., 2024; Forni *et al*., 2013).

**Figure 4.**
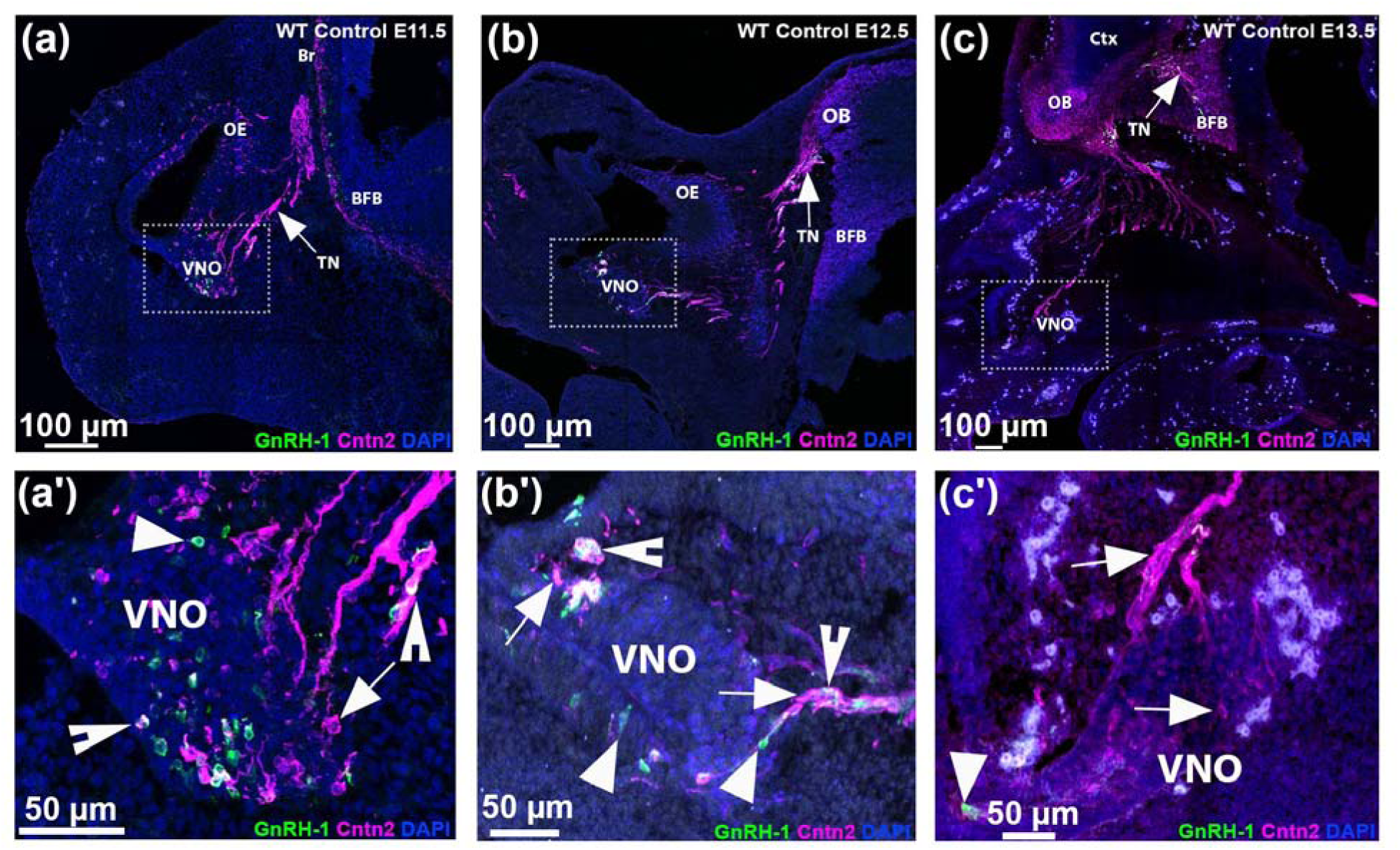
Transient immunoreactivity of Cntn2 in the GnRH-1 neurons within the developing olfactory system. (a-c) developmental time-course of GnRH-1 (green, arrowheads) and Cntn2 (magenta, arrows) immunoreactivity in the developing olfactory system at E11.5, E12, and E13.5. (a-a’) At E11.5, three distinct populations of neurons can be observed, GnRH-1 neurons negative for Cntn2, Cntn2 neurons negative for GnRH-1 and GnRH-1 neurons that express Cntn2 (notched arrowheads). The three distinct populations of neurons are not migrating and are within the vomeronasal organ (VNO). (b-b’) Visualizing at E12.5, the three groups of neurons migrate out of the VNO past the olfactory epithelium (OE) and towards the olfactory bulb (OB). The fibers of the terminal nerve (TN) are immunoreactive for Cntn2. (c-c’) At E13.5, the majority of migratory neurons have left the VNO and have now invaded the brain. The Cntn2+ TN can be observed projecting into the basal forebrain (BFB) Scale bars in a’-c’, 50 µm; a-c, 100 µm.

Based on Cntn2 and GnRH-1 immunoreactivity, we identified three populations: cells only positive for Cntn2, cells reactive for both Cntn2 and GnRH-1, and cells only positive for GnRH1. Specifically, we found that at E11.5, only 15% of the GnRH-1 immuno-positive neurons expressed detectable Cntn2, 47% at E12.5, and 39% at E13.5 (Fig. 4A-C). These data suggest asynchronous and transient expression of Cntn2 across the GnRH-1 neurons.

### Cntn2 can be used as a genetic entry point to manipulate the cells of the nasal migratory mass

Prokr2Cre lineage tracing and Map2 immunoreactivity can be used to visualize the cell bodies of the developing TN (Amato *et al*., 2024).

Previously acquired single-cell RNA sequencing data at E14.4, indicate Cntn2 to be enriched in the TN neurons compared to other neurons in the developing olfactory system, (Fig. 5A) (Amato *et al*., 2024). Based on this, we decided to monitor Cntn2 immunoreactivity in the migratory TN of Prokr2Cre traced animals at various developmental stages. We characterized Cntn2 expression in Prokr2 traced cells using Prokr2Cre^+/-^/Ai14^+/-^ mice at E11.5-E13.5 **(Fig. 5B-D)**.

**Figure 5.**
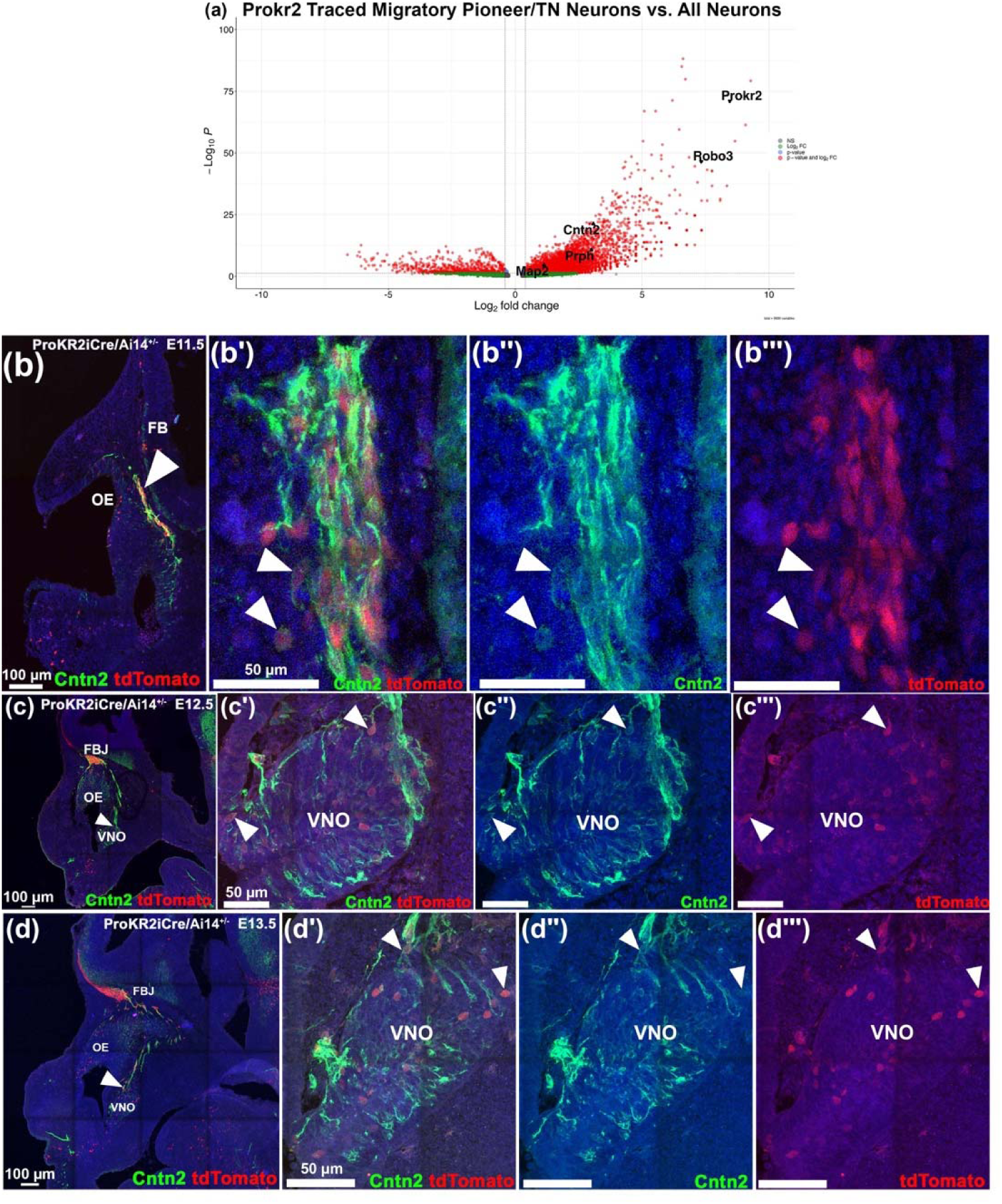
Contactin-2 (Cntn2) as a genetic entry point to manipulate the terminal nerve: (a) Volcano plot comparing enriched genes of the sorted migratory pioneer/TN neurons with all unsorted neurons. (b-d) Developmental time-course of Cntn2 immunoreactivity in the developing vomeronasal organ (VNO) and migratory TN from E11.5-E13.5 using Prokr2Cre^+/-^/Ai14^+/-^ mice. Immunostaining for Cntn2 (green) and Prokr2 lineage tracing (red) was performed at all three developmental stages. (b-b’’’) At E11.5, reveals Cntn2 immunoreactivity in Prokr2 traced pioneer/TN neurons emerging from the olfactory epithelium (OE) proximal to the forebrain (FB). (c-c’’’) Cntn2 immunoreactivity is detected in the traced Prokr2 cells at E12.5, migrating from the VNO to the forebrain junction (FBJ). (d-d’’’) Visualizing at E13.5, an increased number of Prokr2 traced neurons positive for Cntn2 are present in the VNO and FBJ. Scale bars in b’-b’’’, c’-c’’’, d’-d’’’ 50 µm; b-d 100 µm.

At E11.5, Cntn2 immunoreactivity was found in the majority (∼70%) of the Prokr2 traced pioneer/TN neurons emerging from the developing olfactory/vomeronasal epithelium (OE/VNE). However, as Cntn2 has a transient expression, its immunoreactivity dropped to 58% at E12.5 and to 35% of the Prokr2 traced cells at E13.5.

Based on these observations, we identified E11.5 as an attractive time point to induce the Cntn2-Cre-driven genetic tracing in the migratory neurons forming the presumptive TN.

### Using Cntn2^CreERT2^ tracing to follow the pioneer neurons/terminal nerve

Cntn2^CreERT2^ ^+/-^ mice were mated with Ai14^+/-^ reporter mice. Pregnant dams were injected with tamoxifen (75 mg/kg body weight) at E11.5. We analyzed the Cntn2 tracing at E12.5 (1-day post-injection [DPI]), E13.5 (2 DPI), and E15.5 (4 DPI). We combined Cntn2 tracing with Cntn2 immunostaining (Fig. 6A-C). We quantified the percentage of Cntn2-traced cells displaying Cntn2-immunoreactive bodies (Fig. 6D). We found Cntn2 tracing and immunoreactivity overlapping in 15% of the cells at E12.5, 40% at E13.5 and in 20% of the cells at E15.5, providing more evidence for Cntn2’s transient expression pattern. Cntn2 tracing was observed to label many migratory TN cells within the nasal area at E12.5 and E13.5. Notably we observed a larger number of cells over time with a significant increase at E13.5 (Fig. 6E, P=0.0002). However, from E13.5 to E15.5, a significant decrease in the number of traced cell bodies was observed in the nose, suggesting that a portion of the traced neurons might have either died or migrated into the brain (Fig. 6E, P=0.0077). This prompted us to investigate Cntnt2Cre recombination in the GnRH-1 neurons.

**Figure 6.**
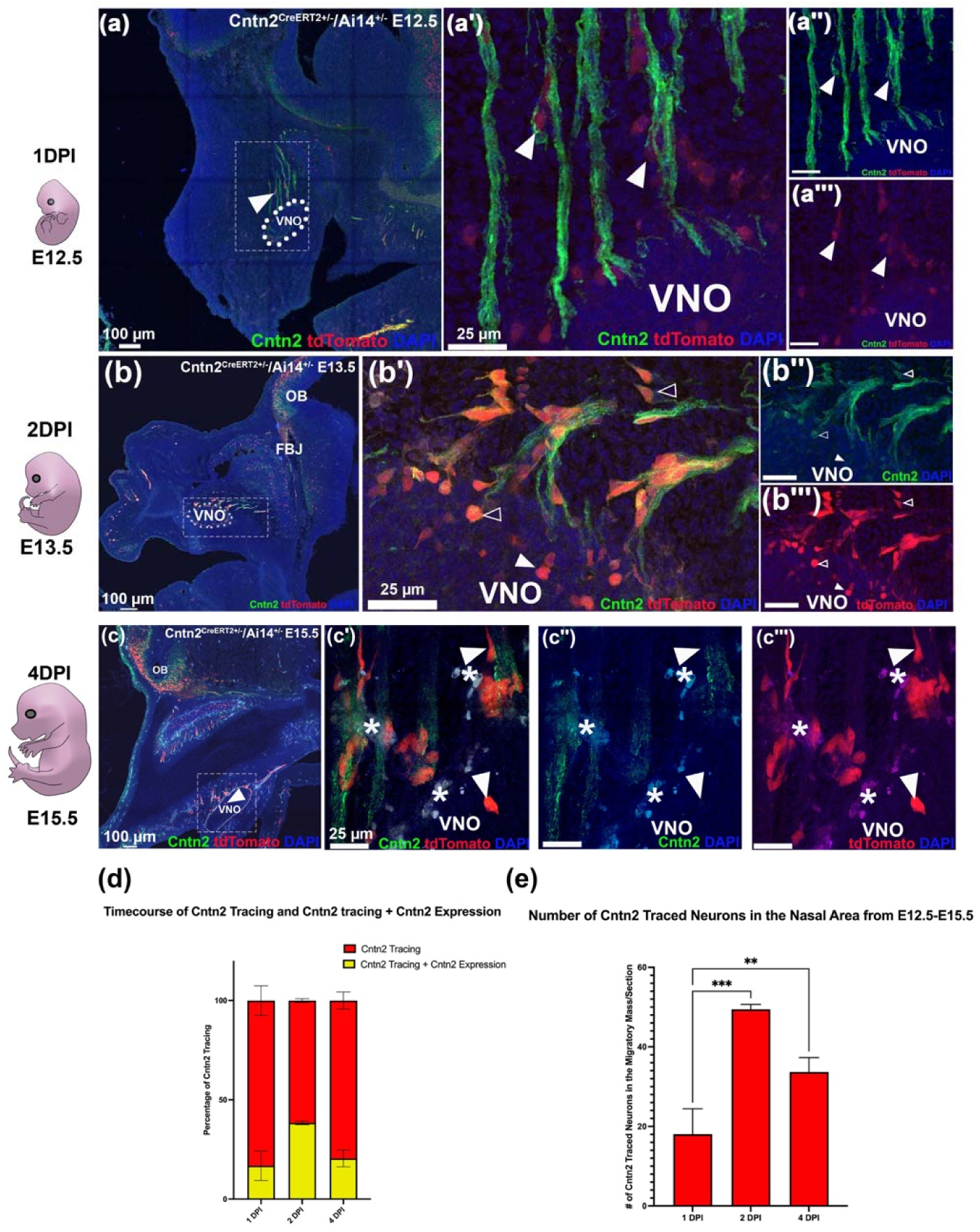
Using Cntn2^CreERT2^ to label cell bodies and Cntn2 to highlight the axons of pioneer neurons/terminal nerve (TN). (a-d) developmental time-course of Cntn2 immunoreactivity in the developing vomeronasal organ (VNO) and migratory TN from E12.5 (1 DPI), E13.5 (2 DPI), and E15.5 (4 DPI) using Cntn2^CreERT2^ ^+/-^/Ai14^+/-^ mice. Immunostaining for Cntn2 (green) and Cntn2 lineage tracing (red) was performed at all three developmental stages. Arrowheads indicate neurons that are traced for Cntn2 and empty arrowheads indicate neurons that are both traced for Cntn2 and expression Cntn2. (a-a’’’) E12.5 reveals Cntn2 traced pioneer/TN neurons labelling cell bodies and Cntn2 immunoreactivity within the axons and fibers. (b-b’’’) At E13.5, Cntn2 immunoreactivity was found to be greater in Cntn2 traced neurons migrating out of the VNO, toward the forebrain junction (FBJ) proximal to the olfactory bulb (OB). (c-c’’’) At E15.5, Cntn2 immunoreactivity was not detected in the Cntn2 traced neurons near the VNO. Red blood cells are marked with asterisks. (d) Quantifications of Cntn2 tracing and traced neurons that positive for Cntn2 expression from E12.5-E15.5, show an increase in Cntn2 expression at E13.5. (e) Quantification of Cntn2 traced neurons in the nasal area from E12.5-E15.5 show significant changes over time (+/- SEM, one-way ANOVA, P>0.05). Scale bars in a’-a’’’, b’-b’’’, c’-c’’’, 25 µm; a, b, c, 100 µm.

### GnRH-1 neurons are positive for Cntn2 Tracing

To understand if a portion of the Cntn2 traced migratory neurons are the GnRH-1 neurons, we performed immunostaining against GnRH-1 on Cntn2^CreERT2+/-^ traced animals at 1,2 and 4 DPI (Tamoxifen injection at E11.5). Both Cntn2 traced neurons and traced neurons also expressing GnRH-1 were observed in the migratory mass together with GnRH-1 neurons negative for tracing (Fig. 7).

**Figure 7.**
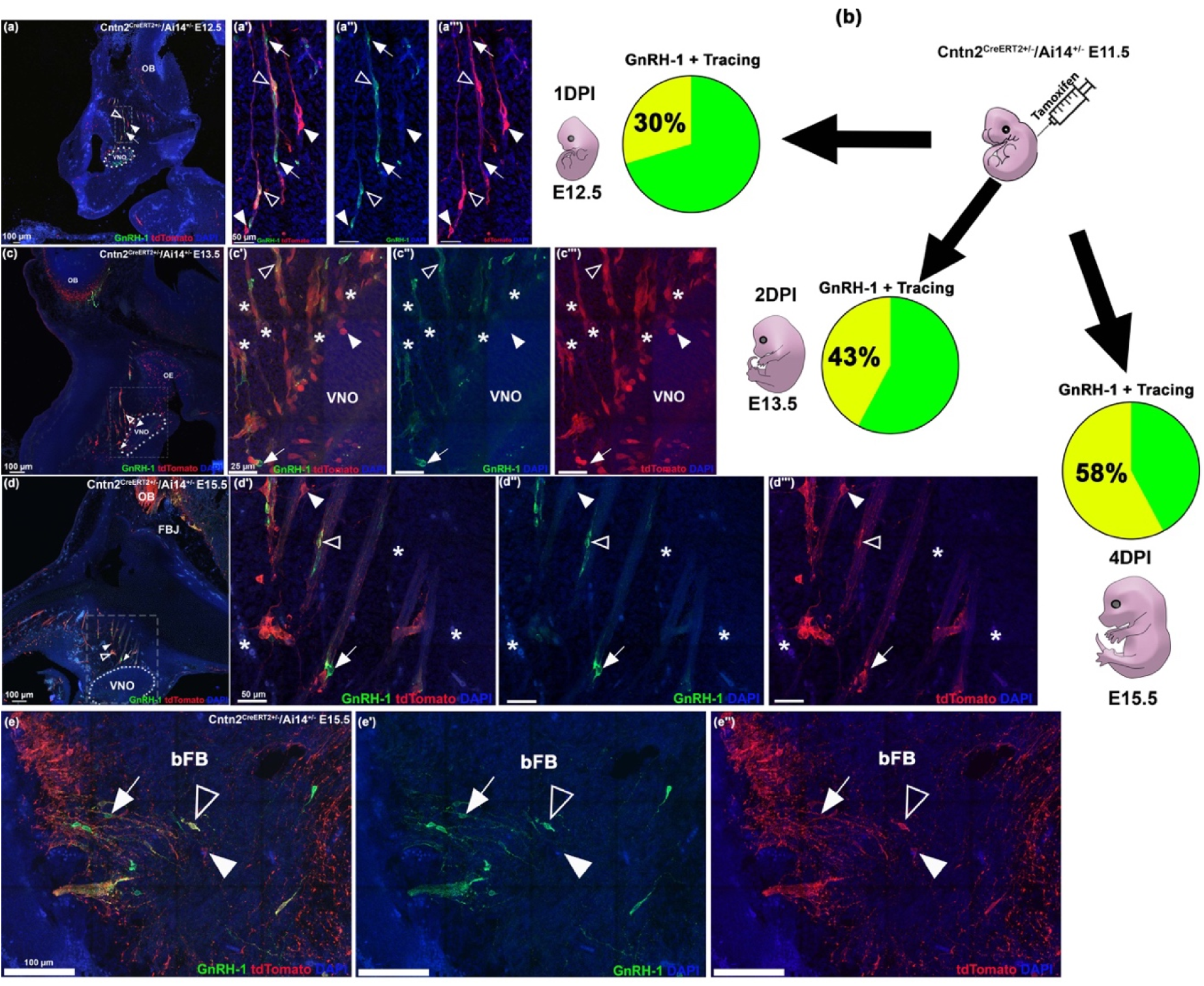
The GnRH-1 neurons are positive for Contactin-2 (Cntn2) tracing: (a-e’’’) Double immunostaining of GnRH-1 (green, arrows) with Cntn2 tracing (red, arrowheads) and GnRH-1 neurons that are traced for Cntn2 (yellow, empty arrowheads) in a time course from E12.5 (1 day post-injection (DPI)), 2 DPI E13.5 and 4 DPI E15.5. (b) Cntn2^CreERT2^ ^+/-^/Ai14^+/-^ mice were injected with tamoxifen at E11.5 and collected at E12.5-E15.5. The relationship between GnRH-1 neurons (green) and GnRH-1 neurons that are traced (yellow) is depicted by the pie charts next to the images. (a-a’’’) E12.5 shows Cntn2 traced neurons, GnRH-1 neurons, and Cntn2 traced neurons positive for GnRH-1 expression within and outside of the vomeronasal organ (VNO); (c-c’’’) At E13.5 there is an increase of the three populations migrating out of the VNO; Cntn2 traced neurons can be seen in the olfactory epithelium (OE) and olfactory bulb (OB). Red blood cells are marked with asterisks. (d-d’’’) E15.5 shows all three populations of neurons invading the brain at the forebrain junction (FBJ), while very few are still migrating out from the VNO. Red blood cells are marked with asterisks. (e-e’’’) Shows a magnification of the basal forebrain (bFB) at E15.5, with the three populations of neurons invading. Scale bars in c’-c’’’, 25 µm; a’-a’’’, d’-d’’’ 50 µm; a, c, d, e, 100 µm.

Interestingly, we observed after one tamoxifen injection at E11.5, the number of traced cells immunoreactive for GnRH-1 increased over time from 30% at 1 DPI to 58% at 4 DPI (Fig. 7A-D), suggesting that Cntn2 expression and therefore recombination precedes GnRH-1 immune detectability. Notably, the Cntn2 recombination in GnRH-1 neurons resulted to be larger than what we expected based on Cntn2 immunoreactivity in the cell bodies at E11.5 (15%). Cntn2 tracing+ TN projections into the basal forebrain (bFB) were observed at E15.5 proximal to GnRH-1 immunoreactive fibers (Fig. 7E).

The GnRH-1 neurons become post-mitotic in a non-synchronous fashion (Jasoni *et al*., 2009). To further evaluate Cntn2 tracing and expression in GnRH-1 neurons, we injected Cntn2^CreERT2^ ^+/-^ animals at E11.5 and E12.5, collecting at E15.5, a stage at which the GnRH-1 neurons invade the brain extensively (Fig.8). From this experimental paradigm, we found a similar number of GnRH-1 neurons were traced (50%), comparable to one injection at E11.5.

**Figure 8.**
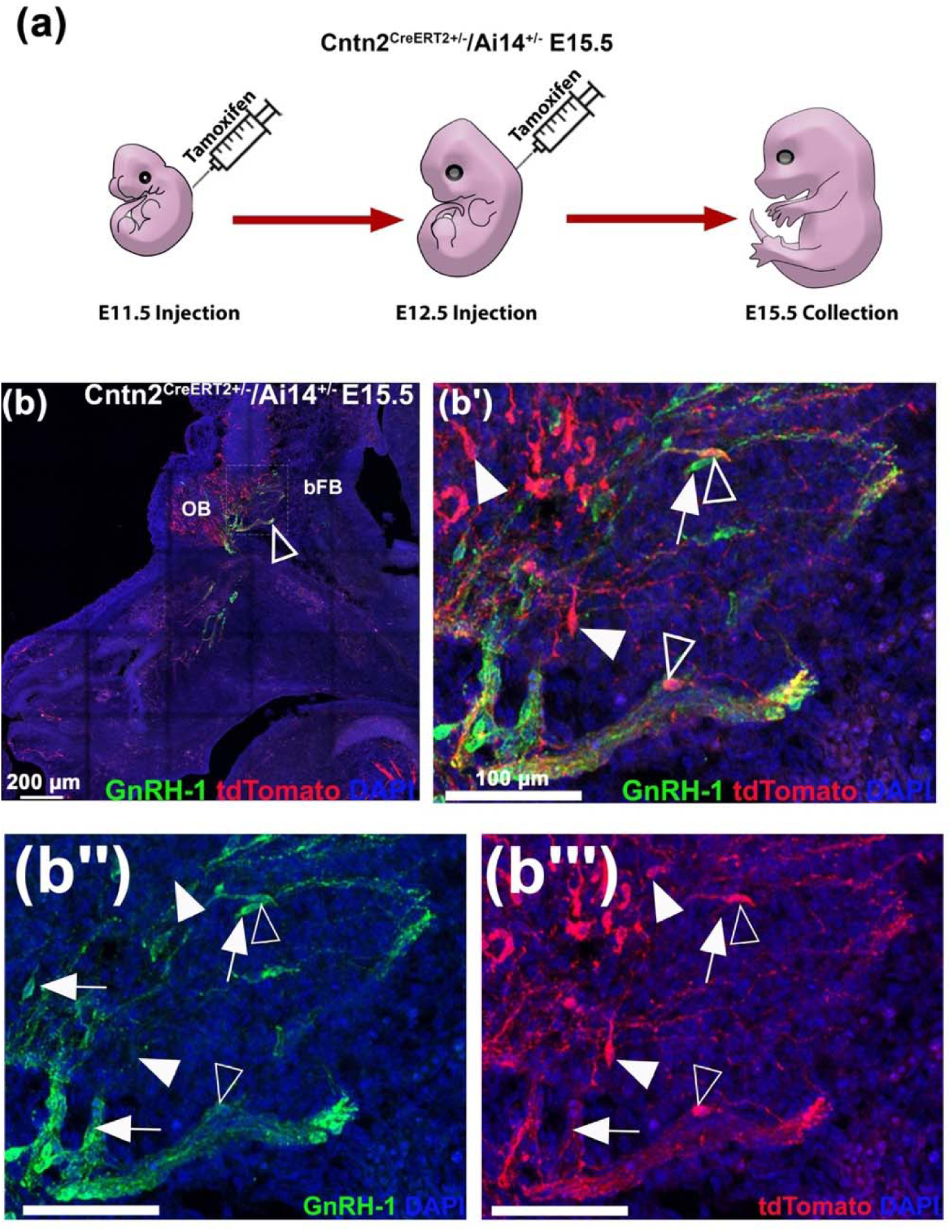
Double injections at E11.5 and E12.5 trace more GnRH-1 neurons. (a) Cntn2^CreERT2^ ^+/-^/Ai14^+/-^ mice were injected with tamoxifen at E11.5 and E12.5 and collected at E15.5. (b-b’’’) Double immunostaining of GnRH-1 (green, arrows) and Cntn2 tracing (red, arrowheads) at E15.5. Many GnRH-1 neurons are observed to be invading the basal forebrain (bFB) with Cntn2 traced neurons and GnRH-1 neurons that are traced for Cntn2 (empty arrowheads). The olfactory bulb (OB) was positive for Cntn2 tracing. Scale bars in b, 200 µm; b’-b’’’, 100 µm.

Notably, we observed traced cell bodies negative for GnRH and TN fibers at the forebrain junction and invading the brain, suggesting that cells of the putative TN might invade the brain together with the GnRH-1 neurons. The transcription factor Isl1 is an early marker for maturing GnRH-1 neurons, which is expressed by virtually all GnRH cells in mice (Taroc *et al*., 2020a; Zouaghi *et al*., 2025). We therefore performed Isl1 immunostaining on Cntn2^CreERT2^ ^+/-^ traced animals. As observed for GnRH-1 immunoreactivity we also found that the percentage of Isl1 immunodetectable traced neurons increases over time: from 22% at 1 DPI to 48% at 2 DPI. This suggests that Cntn2 expression precedes both Isl1 and GnRH-1 immune detectability.

## Discussion

Temporally controlled chimeric genetic recombination in neurons is a valuable method for studying developmental and physiological roles of genes in specific neuronal subsets at distinct developmental stages. During the early stages of the development of the peripheral and central nervous systems, neurons share a large number of generic neuronal genes, making cell type-specific recombination or tracing a challenging task (Fornaro *et al*., 2007).

The olfactory placode is among the very first neurogenic areas in rodent embryos. In fact, migratory neurons can be found emerging from the invaginating olfactory placode as early as E10 which is around two days before the cortical neurogenesis starts (Chen *et al*., 2017).

Contactin-2 can exhibit a transient expression modality in developing neurons, which is why it was previously referred to as transient axonal glycoprotein-1 (TAG1) (Wolfer *et al*., 1994; Yoshida *et al*., 1995). Cntn2’s differential, asynchronous and transient expression across neuronal types, has made Cntn2 a useful epitope to follow specific types of neurons at defined developmental stages.

In this paper, we introduced and characterized a newly generated Cntn2^CreERT2^ mouse line, a novel genetic tool for temporally controlled chimeric recombination in neurons. Our data show that this Cntn2-P2A-CreERT2 model faithfully recombines in Cntn2-expressing neurons, such as the commissural neurons (Fig. 1,2) of the spinal cord, dorsal root ganglia (Dodd *et al*., 1988; Shu *et al*., 2022; Suter *et al*., 2020) as well as in the migratory pioneer/TN neurons of the developing olfactory system, including the GnRH-1 neurons (Figs. 6,7,8) (Ware *et al*., 2016; Wolfer *et al*., 1994; Yamamoto and Schwarting, 1991).

The transient and asynchronous expression of Cntn2 across neurons makes the Cntn2^CreERT2^ mouse line a valuable tool for neurodevelopmental studies aimed at understanding the role of specific guidance molecules or genes involved in neuronal migration (Fig. 3,4).

Cntn2 expression in the developing olfactory area has been observed in neurons believed to form the TN, which serves as the GnRH-1 migratory scaffold (Casoni *et al*., 2016; Duittoz *et al*., 2022; Schwanzel-Fukuda and Pfaff, 1989; Taroc *et al*., 2017). However, given the strong immunoreactivity in the axons and weak immunoreactivity in the cell soma, along with the highly intertwined cell bodies and axons of the olfactory migratory mass, it has never been fully established whether the GnRH-1 neurons themselves express Cntn2.

We recently discovered that Prokr2 Cre recombination selectively labels the putative TN neurons, which we described as migratory neurons of the nasal area that are neither immunoreactive for GnRH-1 nor for the transcription factor Isl1. Notably, Isl1 is a transcription factor that is highly enriched in the GnRH-1 neurons (Lund *et al*., 2020; Taroc *et al*., 2020a; Zouaghi *et al*., 2025). Single-cell RNA sequencing of Prokr2+ presumptive TN cells revealed that Cntn2 is a highly enriched gene in these neurons (Fig. 5) (Amato *et al*., 2024; Martin *et al*., 2011; Pitteloud *et al*., 2007).

Using Prokr2Cre recombination as a reference, we found that at E11.5, Prokr2 tracing, and Cntn2 immunoreactivity overlapped with putative TN cells in the nasal area (70%). Based on this, this stage is appropriate for tamoxifen induction of Cntn2^CreERT2^ recombination in the TN of mice. After Tamoxifen induction at E11.5 the Cntn2^CreERT2+/-^/Ai14^+/-^ embryos were analyzed at multiple intervals post injection (Fig. 6). This analysis revealed recombination in the TN as well as in the GnRH-1 neurons. After a single Tamoxifen injection at E11.5, there was an increase in the number of GnRH-1 neurons positive for immune detectability in traced cells over time (Fig. 7). Isl1 immune detectability was also found to increase in the traced neurons (Fig. S1). This suggests Cntn2 is an early gene expressed prior to GnRH-1/Isl1 immunoreactivity.

Our data suggest that Cntn2 is transiently expressed in both GnRH-1 and TN neurons. Notably at E12.5, we found that approximately half of the GnRH-1 neurons appear to be Cntn2 immunoreactive (Fig. 4). In line with this, Tamoxifen injections in Cntn2^CreERT2+/-^/Ai14^+/-^ at E11.5 and E12.5 traced around 50% of the GnRH-1 neurons (Fig. 8). These data indicate that Cntn2 is a common gene among GnRH-1 neurons and other neurons in the nasal migratory mass and that Cntn2 starts to be expressed before GnRH becomes immunodetectable.

In summary, this study introduces and characterizes the Cntn2^CreERT2^ mouse line. This line serves as a reliable tool for accurately tracing and recombining Cntn2 expressing neurons. By leveraging the transient expression of Cntn2, this model enables precise chimeric tracing of neuronal populations, including commissural neurons, dorsal root ganglia, and migratory neurons in the developing olfactory system.

Analyzing this mouse, we confirmed extensive Cntn2 expression (∼70%) in the Prokr2-traced TN neurons, neurons that we proposed to form the GnRH-1 migratory scaffold. However, tamoxifen-induced recombination at distinct developmental stages highlighted that Cntn2 is also transiently expressed in GnRH-1 neurons, making this model a valuable genetic entry point for studying candidate genes in the early development of the olfactory and GnRH-1 systems.

## Materials and Methods

### Animals

### Mouse lines

The custom mouse line C57BL/6J-Tg(Cntn2-creERT2)^Forni^ was developed using CRISPR/Cas9 technology by Cyagen Biosciences, utilizing the C57BL/6J background strain. Cntn2^CreERT2^ were generated by targeting exon 23 on the Cntn2 gene using CRISPR/Cas9 to insert the donor vector containing the “P2A-CreERT2” cassette (Fig.1A). The targeting vector was designed with homology arms and co-injected into fertilized mouse eggs with the Cas9 mRNA and Cntn2 gRNA (Sequence, matching the reverse strand of the gene: AGCGTTGAGATCAGAGCCTCTGG). F0 Founder animals were identified through PCR and subsequently bred with C57BL/6J wild-type mice to assess germline transmission and facilitate further F1 animal generation. All were meticulously conducted to ensure precise genetic integration.

R26tdTomato Ai14 (JAX stock 007914) mice were maintained on a C57 BL/6J background. Cntn2^CreERT2^ and Ai14 genotypes were confirmed using PCR analysis using the following primers for Ai14: IMR0920: AAGGGAGCTGCAGTGGAGTA; IMR9021: CCGAAAATCTGTGGGAAGTC; *IMR9103:* GGCATTAAAGCAGCGTATCC; IMR9105: CTGTTCCTGTACGGCATGG. For Cntn2^CreERT2^ two primer sets were used interchangeably, generic Cre primers: Cre FWD: AGG TGT AGA GAA GGC ACT TAG C; Cre RVS: CTA ATC GCC ATC TTC CAG CAG G; and specific Cntn2^CreERT2^ primers: Cntn2 WT F4: TAAGGCCTCCATATGACTTTCCTC; Cntn2 WT R7: AAAATTCCTTGCCTGGTTCTATCC; Cntn2 Cre F7: AAGAACGTGGTGCCCCTCTAT; Cntn2 Cre R7: AAAATTCCTTGCCTGGTTCTATCC. Animal euthanasia was done using CO_2_, then cervical dislocation.

Experiments were performed with mice of either sex.

All experiments using animals were completed in accordance with the guidelines of Animal Care and Use Committee at the University at Albany, SUNY.

### TAM preparation and treatment

Tamoxifen (Sigma–Aldrich), CAS # 10540−29−1, was mixed and dissolved in corn oil at a concentration of 20 µg/µl. Time-mated females were injected intraperitoneally at the chosen embryonic day with a dosage of 75 mg/Kg body weight, and embryos were collected at the desired time point. The observation of the vaginal plug from time-mated females was defined as E0.5 to determine embryo age.

### Tissue Preparation

Collected embryos were fixed in 3.7% formaldehyde/PBS at 4°C for 2 h, then submerged in 30% sucrose overnight. Embryos were embedded and frozen in O.C.T. (Tissue-TeK) and stored at −80°C. Cryosectioning was done using a CM3050S Leica cryostat, samples were collected on Super-frost plus slides (VWR) at 18 µm thickness for immunofluorescent staining.

### Immunohistochemistry

#### Immunofluorescence

Primary antibodies and dilutions used for this study: goat-α-Cntn2 (1:1000, R&D Systems), chicken-α-RFP (1:1000, Rockland), SW rabbit-α-GnRH-1 (1:6000, Susan Wray, NIH), mouse-α-Isl1 (1:100, DSHB) and rabbit-α-RFP (1:500, Rockland). Experiments using mouse-α-Isl1 and rabbit-α-RFP required antigen retrieval which was done by submerging slides in citric acid solution heated to 95°C for 15 min, then cooling for 15 minutes before proceeding with the standard immunofluorescence protocol. Secondary antibodies matched for the correct species were conjugated to Alexa Fluor-488, Alexa Flour-594, or Alexa Flour-680 (Invitrogen and Jackson Laboratories). Counterstaining was done using 4′,6′-diamidino-2-phenylindole (1:3000;Sigma-Aldrich) and coverslips were mounted with Fluorogel l (Electron Microscopy Sciences). Confocal microscopy image acquisition was taken with a LSM 980 microscope (Zeiss). FIJI/ImageJ was used for image analysis and quantifications. Each staining presented was replicated on three different animals.

#### Chromogen-based reactions

Experiments using chromogen-based reactions were stained as described previously (Forni *et al*., 2011a). The tissue was processed using a standard avidin–biotin– horseradish peroxidase/3,3-diaminobenzidine (DAB) procedure. Each staining was visualized using nickel (II) sulfate heptahydrate (Sigma) to intensify the DAB reaction (black) followed by a counterstain with methyl green. Brightfield images were taken on a Leica DM4000 B LED fluorescence microscope equipped with a Leica DFC310 FX 422 camera. FIJI/ImageJ software was used for further evaluation. Staining presented were replicated on three different animals.

### Cell quantifications

All quantifications were performed using the cell counter plugin within FIJI/ImageJ. Quantification of Cntn2 tracing and Cntn2 immunoreactivity in the spinal cord was performed on E12.5 Cntn2^CreERT2^ traced transverse sections, and the cell bodies of the dorsal root ganglia were quantified. For all quantifications in the developing nasal area, E12.5-E15.5 Cntn2^CreERT2^ traced parasagittal sections were used. For each immunostaining (Cntn2, GnRH-1 and Isl1) paired with Cntn2 tracing, cell bodies in the nasal area/migratory mass were quantified. For all quantifications, the number of cells per section (∼6 sections) was averaged for each animal.

### Statistics

Statistical analyses were conducted using Graphpad Prism 10.4.1. At least three biological replicates were used when performing each quantification. Significance was determined to be p value < 0.05 and was assessed using a one-way ANOVA or two-way ANOVA where applicable.

### Single-Cell RNA Sequencing

The volcano plot generated using single-cell sequencing data available through the Gene Expression Omnibus (GEO) database, under GEO accession number GSE234871 (https://www.ncbi.nlm.nih.gov/geo/query/acc.cgi?acc=GSE234871). These were previously described (Amato *et al*., 2024).

## Acknowledgments

This work was supported by the Eunice Kennedy Shriver National Institute of Child Health and Human Development (NICHD) under Grants 2R01HD097331 (P.E.F.) and 1R01HD114827 (P.E.F.), as well as by the National Institute on Deafness and Other Communication Disorders (NIDCD) under Grant R01DC017149 (P.E.F.). The Zeiss 980 microscope at the University at Albany was funded by the Office of the Director, NIH, under Award Number S10OD028600.

---

**Figure S1.**
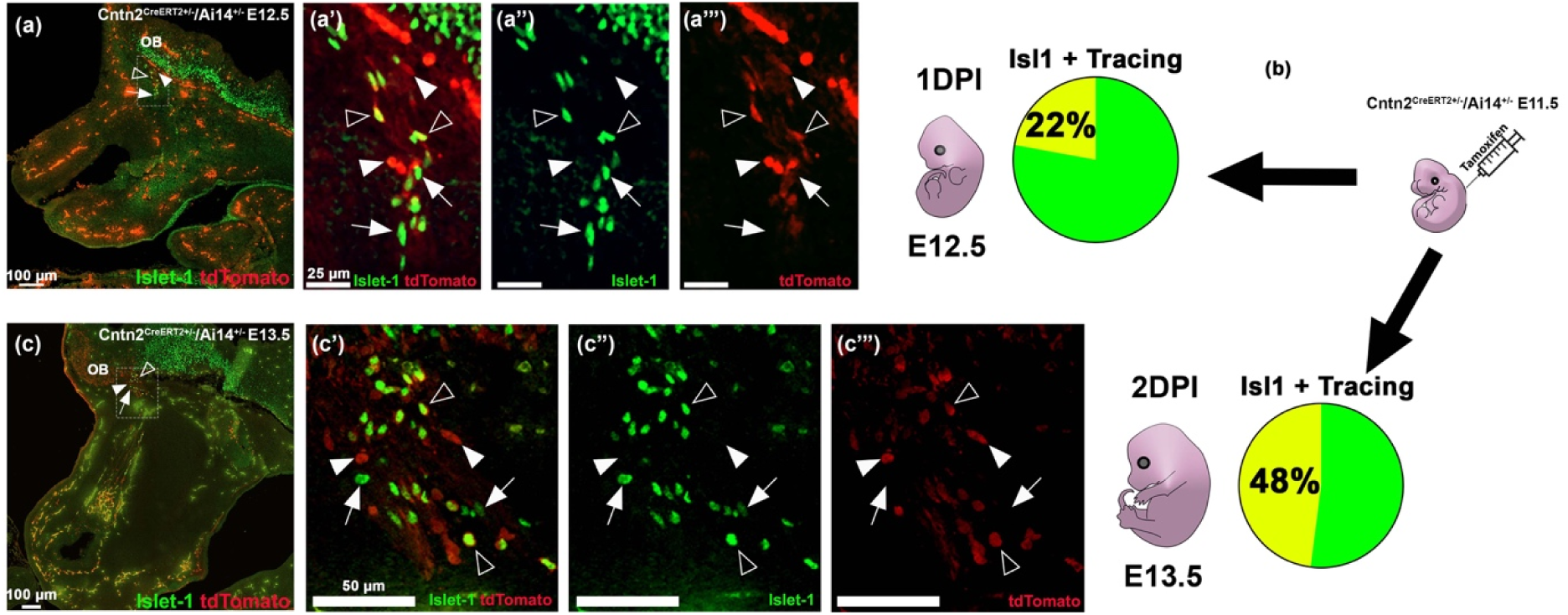
Isl1 immunoreactivity precedes Cntn2 expression. (a-c) Double immunofluorescent staining of Isl1 (green, arrows) and Cntn2 tracing (red, arrowheads). (b) Cntn2^CreERT2^ ^+/-^/Ai14^+/-^ mice were injected with tamoxifen at E11.5 and collected at E12.5 and E13.5. Isl1 expression is depicted by the pie charts next to the images, Isl1+ neurons (green) and Isl1 neurons that are traced (yellow). (a-a’’’, c-c’’’) At E12.5 and E13.5, migratory neurons expressing Isl1, Cntn2 tracing, and Isl1 immunopositive traced neurons (yellow, empty arrowheads) were observed to be approaching the brain, proximal to the olfactory bulb (OB). Scale bars in a’-a’’’, 25 µm; c’-c’’’, 50 µm; a, c, 100 µm.

## References

Amato E, Jr., Taroc EZM, Forni PE. 2024. Illuminating the terminal nerve: Uncovering the link between GnRH-1 neuron and olfactory development. J Comp Neurol 532: e25599.

Barraud P, Seferiadis AA, Tyson LD, Zwart MF, Szabo-Rogers HL, Ruhrberg C, Liu KJ, Baker CV. 2010. Neural crest origin of olfactory ensheathing glia. Proc Natl Acad Sci U S A 107: 21040–21045.

Barraud P, St John JA, Stolt CC, Wegner M, Baker CV. 2013. Olfactory ensheathing glia are required for embryonic olfactory axon targeting and the migration of gonadotropin-releasing hormone neurons. Biol Open 2: 750–759.

Casoni F, Malone SA, Belle M, Luzzati F, Collier F, Allet C, Hrabovszky E, Rasika S, Prevot V, Chedotal A, Giacobini P. 2016. Development of the neurons controlling fertility in humans: new insights from 3D imaging and transparent fetal brains. Development 143: 3969–3981.

Chao DL, Ma L, Shen K. 2009. Transient cell-cell interactions in neural circuit formation. Nat Rev Neurosci 10: 262–271.

Chen VS, Morrison JP, Southwell MF, Foley JF, Bolon B, Elmore SA. 2017. Histology Atlas of the Developing Prenatal and Postnatal Mouse Central Nervous System, with Emphasis on Prenatal Days E7.5 to E18.5. Toxicol Pathol 45: 705–744.

Demski LS, Schwanzel-Fukuda M. 1987. The terminal nerve (nervus terminalis): structure, function, and evolution. Introduction. Ann N Y Acad Sci 519: ix–xi.

Dodd J, Morton SB, Karagogeos D, Yamamoto M, Jessell TM. 1988. Spatial regulation of axonal glycoprotein expression on subsets of embryonic spinal neurons. Neuron 1: 105–116.

Donnelly MLL, Hughes LE, Luke G, Mendoza H, Ten Dam E, Gani D, Ryan MD. 2001a. The ’cleavage’ activities of foot-and-mouth disease virus 2A site-directed mutants and naturally occurring ’2A-like’ sequences. J Gen Virol 82: 1027–1041.

Donnelly MLL, Luke G, Mehrotra A, Li X, Hughes LE, Gani D, Ryan MD. 2001b. Analysis of the aphthovirus 2A/2B polyprotein ’cleavage’ mechanism indicates not a proteolytic reaction, but a novel translational effect: a putative ribosomal ’skip’. J Gen Virol 82: 1013–1025.

Duittoz AH, Forni PE, Giacobini P, Golan M, Mollard P, Negron AL, Radovick S, Wray S. 2021. Development of the gonadotropin-releasing hormone system. J Neuroendocrinol: e13087.

Duittoz AH, Forni PE, Giacobini P, Golan M, Mollard P, Negron AL, Radovick S, Wray S. 2022. Development of the gonadotropin-releasing hormone system. J Neuroendocrinol 34: e13087.

Fan S, Liu J, Chofflet N, Bailey AO, Russell WK, Zhang Z, Takahashi H, Ren G, Rudenko G. 2024. Molecular mechanism of contactin 2 homophilic interaction. Structure 32: 1652–1666 e1658.

Fornaro M, Geuna S, Fasolo A, Giacobini-Robecchi MG. 2003. HuC/D confocal imaging points to olfactory migratory cells as the first cell population that expresses a post-mitotic neuronal phenotype in the chick embryo. Neuroscience 122: 123–128.

Fornaro M, Raimondo S, Lee JM, Giacobini-Robecchi MG. 2007. Neuron-specific Hu proteins sub-cellular localization in primary sensory neurons. Ann Anat 189: 223–228.

Forni PE, Bharti K, Flannery EM, Shimogori T, Wray S. 2013. The indirect role of fibroblast growth factor-8 in defining neurogenic niches of the olfactory/GnRH systems. J Neurosci 33: 19620–19634.

Forni PE, Fornaro M, Guenette S, Wray S. 2011a. A role for FE65 in controlling GnRH-1 neurogenesis. J Neurosci 31: 480–491.

Forni PE, Taylor-Burds C, Melvin VS, Williams T, Wray S. 2011b. Neural crest and ectodermal cells intermix in the nasal placode to give rise to GnRH-1 neurons, sensory neurons, and olfactory ensheathing cells. J Neurosci 31: 6915–6927.

Furley AJ, Morton SB, Manalo D, Karagogeos D, Dodd J, Jessell TM. 1990. The axonal glycoprotein TAG-1 is an immunoglobulin superfamily member with neurite outgrowth-promoting activity. Cell 61: 157–170.

Gong Q, Shipley MT. 1995. Evidence that pioneer olfactory axons regulate telencephalon cell cycle kinetics to induce the formation of the olfactory bulb. Neuron 14: 91–101.

Gurung S, Asante E, Hummel D, Williams A, Feldman-Schultz O, Halloran MC, Sittaramane V, Chandrasekhar A. 2018. Distinct roles for the cell adhesion molecule Contactin2 in the development and function of neural circuits in zebrafish. Mech Dev 152: 1–12.

Jasoni CL, Porteous RW, Herbison AE. 2009. Anatomical location of mature GnRH neurons corresponds with their birthdate in the developing mouse. Dev Dyn 238: 524–531.

Jennes L. 1992. Selective expression of peripherin in gonadotropin-releasing hormone-synthesizing neurons of the rat. Mol Cell Neurosci 3: 571–577.

Kaprara A, Huhtaniemi IT. 2018. The hypothalamus-pituitary-gonad axis: Tales of mice and men. Metabolism 86: 3–17.

Karagogeos D, Morton SB, Casano F, Dodd J, Jessell TM. 1991. Developmental expression of the axonal glycoprotein TAG-1: differential regulation by central and peripheral neurons in vitro. Development 112: 51–67.

Lund C, Yellapragada V, Vuoristo S, Balboa D, Trova S, Allet C, Eskici N, Pulli K, Giacobini P, Tuuri T, Raivio T. 2020. Characterization of the human GnRH neuron developmental transcriptome using a GNRH1-TdTomato reporter line in human pluripotent stem cells. Dis Model Mech 13.

Martin C, Balasubramanian R, Dwyer AA, Au MG, Sidis Y, Kaiser UB, Seminara SB, Pitteloud N, Zhou QY, Crowley WF, Jr. 2011. The role of the prokineticin 2 pathway in human reproduction: evidence from the study of human and murine gene mutations. Endocr Rev 32: 225–246.

Pitteloud N, Zhang C, Pignatelli D, Li JD, Raivio T, Cole LW, Plummer L, Jacobson-Dickman EE, Mellon PL, Zhou QY, Crowley WF, Jr. 2007. Loss-of-function mutation in the prokineticin 2 gene causes Kallmann syndrome and normosmic idiopathic hypogonadotropic hypogonadism. Proceedings of the National Academy of Sciences of the United States of America 104: 17447–17452.

Pohl CR, Knobil E. 1982. The role of the central nervous system in the control of ovarian function in higher primates. Annu Rev Physiol 44: 583–593.

Schwanzel-Fukuda M, Bick D, Pfaff DW. 1989. Luteinizing hormone-releasing hormone (LHRH)-expressing cells do not migrate normally in an inherited hypogonadal (Kallmann) syndrome. Brain Res Mol Brain Res 6: 311–326.

Schwanzel-Fukuda M, Garcia MS, Morrell JI, Pfaff DW. 1987. Distribution of luteinizing hormone-releasing hormone in the nervus terminalis and brain of the mouse detected by immunocytochemistry. The Journal of comparative neurology 255: 231–244.

Schwanzel-Fukuda M, Pfaff DW. 1989. Origin of luteinizing hormone-releasing hormone neurons. Nature 338: 161–164.

Schwarting GA, Kostek C, Bless EP, Ahmad N, Tobet SA. 2001. Deleted in colorectal cancer (DCC) regulates the migration of luteinizing hormone-releasing hormone neurons to the basal forebrain. J Neurosci 21: 911–919.

Schwarting GA, Raitcheva D, Bless EP, Ackerman SL, Tobet S. 2004. Netrin 1-mediated chemoattraction regulates the migratory pathway of LHRH neurons. Eur J Neurosci 19: 11–20.

Schwarting GA, Wierman ME, Tobet SA. 2007. Gonadotropin-releasing hormone neuronal migration. Semin Reprod Med 25: 305–312.

Shu M, Hong D, Lin H, Zhang J, Luo Z, Du Y, Sun Z, Yin M, Yin Y, Liu L, Bao S, Liu Z, Lu F, Huang J, Dai J. 2022. Single-cell chromatin accessibility identifies enhancer networks driving gene expression during spinal cord development in mouse. Dev Cell 57: 2761–2775 e2766.

Suter T, Blagburn SV, Fisher SE, Anderson-Keightly HM, D’Elia KP, Jaworski A. 2020. TAG-1 Multifunctionality Coordinates Neuronal Migration, Axon Guidance, and Fasciculation. Cell Rep 30: 1164–1177 e1167.

Suter T, Jaworski A. 2019. Cell migration and axon guidance at the border between central and peripheral nervous system. Science 365.

Tang W, Ehrlich I, Wolff SB, Michalski AM, Wolfl S, Hasan MT, Luthi A, Sprengel R. 2009. Faithful expression of multiple proteins via 2A-peptide self-processing: a versatile and reliable method for manipulating brain circuits. J Neurosci 29: 8621–8629.

Taroc EZM, Katreddi RR, Forni PE. 2020a. Identifying Isl1 Genetic Lineage in the Developing Olfactory System and in GnRH-1 Neurons. Front Physiol 11: 601923.

Taroc EZM, Lin JM, Tulloch AJ, Jaworski A, Forni PE. 2019. GnRH-1 Neural Migration From the Nose to the Brain Is Independent From Slit2, Robo3 and NELL2 Signaling. Front Cell Neurosci 13: 70.

Taroc EZM, Naik AS, Lin JM, Peterson NB, Keefe DL, Jr., Genis E, Fuchs G, Balasubramanian R, Forni PE. 2020b. Gli3 Regulates Vomeronasal Neurogenesis, Olfactory Ensheathing Cell Formation, and GnRH-1 Neuronal Migration. J Neurosci 40: 311–326.

Taroc EZM, Prasad A, Lin JM, Forni PE. 2017. The terminal nerve plays a prominent role in GnRH-1 neuronal migration independent from proper olfactory and vomeronasal connections to the olfactory bulbs. Biol Open 6: 1552–1568.

Tessier-Lavigne M, Placzek M, Lumsden AG, Dodd J, Jessell TM. 1988. Chemotropic guidance of developing axons in the mammalian central nervous system. Nature 336: 775–778.

Ware M, Hamdi-Roze H, Le Friec J, David V, Dupe V. 2016. Regulation of downstream neuronal genes by proneural transcription factors during initial neurogenesis in the vertebrate brain. Neural Dev 11: 22.

Wolfer DP, Henehan-Beatty A, Stoeckli ET, Sonderegger P, Lipp HP. 1994. Distribution of TAG-1/axonin-1 in fibre tracts and migratory streams of the developing mouse nervous system. J Comp Neurol 345: 1–32.

Wolman MA, Sittaramane VK, Essner JJ, Yost HJ, Chandrasekhar A, Halloran MC. 2008. Transient axonal glycoprotein-1 (TAG-1) and laminin-alpha1 regulate dynamic growth cone behaviors and initial axon direction in vivo. Neural Dev 3: 6.

Wray S. 2001. Development of luteinizing hormone releasing hormone neurones. J Neuroendocrinol 13: 3–11.

Wray S, Hoffman G. 1986. A developmental study of the quantitative distribution of LHRH neurons within the central nervous system of postnatal male and female rats. J Comp Neurol 252: 522–531.

Wray S, Key S, Qualls R, Fueshko SM. 1994. A subset of peripherin positive olfactory axons delineates the luteinizing hormone releasing hormone neuronal migratory pathway in developing mouse. Dev Biol 166: 349–354.

Wray S, Nieburgs A, Elkabes S. 1989. Spatiotemporal cell expression of luteinizing hormone-releasing hormone in the prenatal mouse: evidence for an embryonic origin in the olfactory placode. Brain Res Dev Brain Res 46: 309–318.

Yamamoto M, Boyer AM, Crandall JE, Edwards M, Tanaka H. 1986. Distribution of stage-specific neurite-associated proteins in the developing murine nervous system recognized by a monoclonal antibody. J Neurosci 6: 3576–3594.

Yamamoto M, Schwarting G. 1991. The formation of axonal pathways in developing cranial nerves. Neurosci Res 11: 229–260.

Yoshida K, Tobet SA, Crandall JE, Jimenez TP, Schwarting GA. 1995. The migration of luteinizing hormone-releasing hormone neurons in the developing rat is associated with a transient, caudal projection of the vomeronasal nerve. J Neurosci 15: 7769–7777.

Zouaghi Y, Alpern D, Gardeux V, Russeil J, Deplancke B, Santoni F, Pitteloud N, Messina A. 2025. Transcriptomic profiling of murine GnRH neurons reveals developmental trajectories linked to human reproduction and infertility. Theranostics 15: 3673–3692.

